# Fitness effects of local seasonal adaptation depend on the timing of reproduction in a range-expanding butterfly

**DOI:** 10.64898/2026.01.28.702196

**Authors:** Martina Bovolenta, Mats Ittonen, Karl Gotthard

## Abstract

Local seasonal adaptation across latitudes is ubiquitous, but its fitness consequences are rarely estimated in the field. Using a common-garden field experiment at the northern range margin of the butterfly *Lasiommata megera*, we show that fitness consequences of among-population, genetic differences in photoperiodism for diapause (dormancy) timing depend on when during the adult flight period eggs are laid. In early-developing cohorts, individuals from southern populations were more likely to avert diapause than locally adapted northern range margin populations, and this non-diapause development was strongly disfavoured by natural selection. However, for all populations, virtually all eggs laid only one week later entered diapause, limiting overall among-population phenotypic differences. This demonstrates how evolved genetic differences for a reaction norm interact with phenology to shape the developmental decisions of caterpillars in the wild. Rapid local adaptation of photoperiodism has likely occurred through natural selection acting on a limited part of each autumn generation, implying a smooth fitness landscape where mildly locally maladapted populations can establish and subsequently evolve towards the local fitness peak. Thus, latitudinal differences in daylength are unlikely to restrict climate change–driven range expansions.

## 1. Introduction

Climate change lengthens growing seasons and makes previously too cold areas suitable for new species, leading to advances in phenology (yearly seasonal timing of life cycle events), increases in voltinism (number of generations per year), and poleward range expansions (Parmesan & Yohe, 2003; Altermatt, 2010; Chen *et al*., 2011; Pöyry *et al*., 2011; Mason *et al*., 2015; Cohen *et al*., 2018). In addition to direct effects on populations, the alteration of seasonal environments can impose novel selection pressures on existing adaptations for life cycle timing. In many organisms seasonal timing is regulated by daylength – a calendar unaffected by short-term fluctuations in weather (Nelson *et al*. 2010) – but because photoperiod does not change with climate, this otherwise reliable cue can become mismatched with shifting seasonal conditions in a warming world (Forrest, 2016). As a result, developmental decisions may occur at the wrong time of year. Altered seasonal timing (e.g. advanced phenology) or faster growth can shift daylength-sensitive life stages into different photoperiodic environments, and poleward range expansions expose populations to novel photoperiodic regimes, potentially limiting the ability of range-shifting organisms to track suitable climates (Clausen & Clausen, 2013; Grevstad & Coop, 2015; Van Dyck *et al*., 2015; Way & Montgomery, 2015). However, such novel selection pressures are likely to be important only during a limited part of the life cycle. Because a longer growth season comes about through an earlier start and/or a later end of the favourable season, selection will be particularly strong on adaptations that regulate entry into and exit from the active part of the life cycle.

Insects in temperate climates usually overwinter in diapause, a state of slowed development and reduced metabolism that allows synchronizing development and reproduction with resource availability (Tauber *et al*., 1986; Denlinger, 2022). Diapause is induced in advance of the onset of adverse conditions, as insects in a sensitive stage respond to cues that signal future conditions. Winter diapause can occur either obligately in each generation or facultatively through plastic responses to environmental cues (most commonly short autumn daylengths), and this facultative diapause is the main adaptation enabling temperate insects to produce multiple generations within a single growing season. The timing of entry into facultative diapause is crucial since most insects can enter diapause only in one life stage (Denlinger, 2022). For example, if an individual of a species with larval diapause pupates late in autumn, it can no longer enter diapause, and its offspring may not have time to reach the diapausing larval stage before winter. Climate change has been proposed to lead to such mistiming – “lost generations” (Van Dyck *et al*. 2015) – in two main ways. First, earlier phenology due to earlier springs and faster development in warm temperatures can make individuals reach their daylength-sensitive life stage earlier in the year, when days are too long to induce diapause. The resulting additional late-season generation is a common response to climate change (Altermatt, 2010) and can be adaptive (Wepprich *et al*., 2025), but may also have severe population-level consequences if the growing season remains too short to allow successful overwintering of the last generation’s offspring (Van Dyck *et al*., 2015). Second, poleward-expanding populations encounter longer daylengths than they have previously evolved to time their life cycles in, potentially leading to an extra adult generation whose offspring, again, become a lost generation (Grevstad & Coop, 2015). This could restrict climate change-driven poleward range expansions.

The lost generation phenomenon is expected to impose strong selection on photoperiodic diapause induction thresholds. These thresholds are generally considered to be highly evolvable (Bradshaw & Holzapfel, 2007; Tanaka & Murata, 2016; Gotthard *et al*., 2026), and latitudinally range-expanding populations have rapidly evolved to better match local photoperiodic cues (Urbanski *et al*., 2012; Bean *et al*., 2012; Ittonen *et al*., 2022). However, studies have rarely demonstrated actual fitness consequences of these putative local adaptations in the field (but see Bradshaw *et al*., 2004), leaving the importance of evolution in response to the cue-environment decoupling unclear. Furthermore, studies that have examined contemporary evolution of diapause induction responses within wild populations have produced mixed results (Nielsen *et al*., 2023; Kaiser & Van Dyck, 2025; Nechols *et al*., 2025; Gomi, 2025), suggesting that selection pressures or evolutionary capacity vary among populations. Explicit tests of the fitness consequences of partial voltinism shifts are needed for determining whether limited evolutionary responses in some systems reflect weak selection or insufficient genetic variation.

The wall brown butterfly, *Lasiommata megera* (Linnaeus 1767), has expanded northwards in Sweden since around the year 2000 (Ittonen *et al*., 2022). Conversely, the species has declined in north-western Europe (Fox *et al*., 2023; Van Dyck *et al*., 2009; van Swaay *et al*., 2025), and lost generations have been proposed to contribute to this decline (Van Dyck *et al*., 2015). A recent study in Belgium found no evidence for evolutionary change mitigating lost generations (Kaiser & Van Dyck, 2025), but along a latitudinal gradient in Sweden, northern range margin populations have evolved to both enter diapause in longer days (Ittonen *et al*., 2022) and grow faster before diapause (Ittonen *et al*., 2025) than southern-Swedish populations. These differences were interpreted as local adaptations: longer-day diapause induction reduces the risk of maladaptive non-diapause development in autumn, and faster growth increases chances of reaching sufficient body mass before winter despite a short growing season. However, in their reciprocal field transplant experiment, Ittonen *et al*. (2025) did not detect fitness consequences of these differences. The authors hypothesised that fitness effects manifest only during specific parts of the autumn generation, uncaptured in their experiment. Differences in diapause induction would primarily affect the earliest-emerging individuals of the last overwintering generation (which face longer daylengths), and growth rate differences would matter mainly for the latest individuals (which have the least time to grow before winter). This hypothesis has important implications for colonization dynamics. When genotypes adapted to southern seasonal conditions expand northwards, most individuals may successfully enter diapause and reach a suitable life stage for overwintering, while maladaptation affects only the earliest and latest individuals within each autumn generation. If so, strong natural selection acting on a limited fraction of each autumn generation has been enough to produce rapid evolution during northward expansion in *L. megera*, and potentially many other insects.

We tested whether the fitness effects of known, among-population, genetic differences in daylength thresholds for diapause induction and growth rate depend on the seasonal timing of oviposition. We performed a common-garden field experiment near the species’ northern range margin in Sweden, placing eggs of butterflies from northern and southern populations into field cages, thereby simulating a long-range dispersal event. We manipulated oviposition timing by introducing eggs at three time points one week apart (creating early-, intermediate-, and late-season cohorts). In late autumn, we assessed diapause induction and, after weighing, allowed diapausing caterpillars to overwinter, thus testing whether population and prewinter mass affect winter survival. Further, we let individuals that did not enter diapause, but instead developed into adult butterflies, mate and lay eggs to determine whether the resulting additional generation could reach the diapausing life stage and survive until spring (or whether it constituted a lost generation). We predicted that (1) individuals of northern descent enter diapause more frequently than southern individuals but that this difference is reduced (or disappears entirely) in the later cohorts that experience daylengths short enough to induce diapause in individuals of all genetic backgrounds; (2) individuals failing to enter diapause would be unable to produce successfully overwintering offspring; (3) northern-descent individuals would grow faster than southern individuals during development into diapause; and (4) higher pre-winter mass would increase overwinter survival, but only in the latest-laid cohort, in which time constraints generate many small individuals.

## 2. Materials and Methods

### 2.1. Study species and population sampling

The wall brown butterfly, *Lasiommata megera* (Nymphalidae: Satyrinae), occurs in a large part of Europe, the northernmost range margin being in Fennoscandia (Ittonen *et al*., 2022). It lives in grass-dominated habitats, such as seashores and dry meadows, and caterpillars feed on various common grasses of the family Poaceae (Wickman, 1988; Frohawk, 1924; Emmet & Heat, 1989). In Sweden, *L. megera* has at least two adult generations per year (bivoltinism), even at the northern range margin, and a partial third generation occurs in warm years, especially in the south (Fig. 1b). Winter diapause typically occurs in the third larval instar; however, caterpillars are sensitive to daylength and modifying temperature cues for diapause induction at least during their second and third instars (Frohawk, 1924; Ittonen *et al*., 2022).

**Figure 1.**
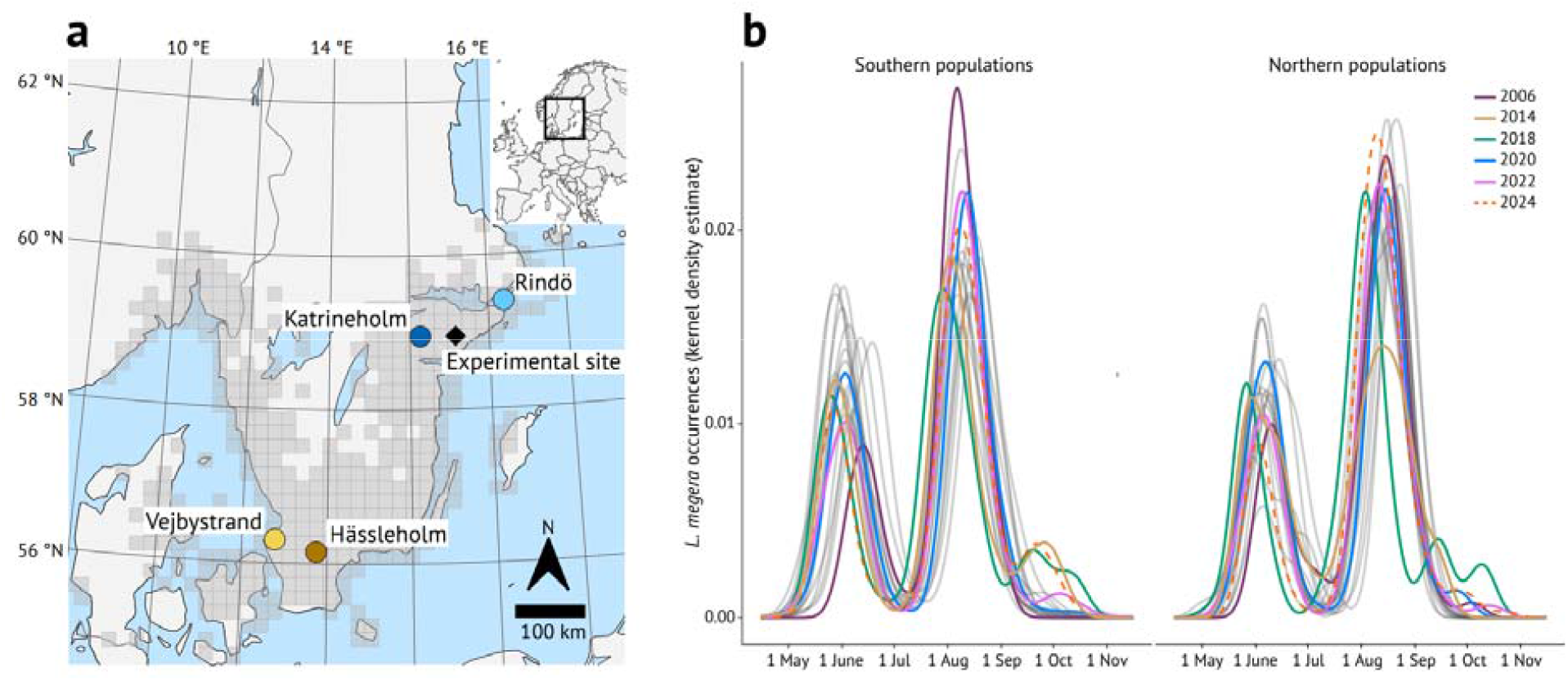
Study populations and the phenology of *Lasiommata megera*. (a) Map of the study locations. Each circle shows a population, and the diamond shows Tovetorp Zoological Station, where the experiment was conducted. The grey shading shows the range of *L. megera* in this part of Europe. (b) The natural phenology of *L. megera* in its southern (south of 58.5 °N; left panel) and northern (north of 58.5 °N; right panel) range in Sweden based on observation data from the years 2006–2025 (GBIF.org, 2026). Note that warm summers (such as the 2018 European heatwave) have led to earlier phenology and a small third generation also at the northern range margin; such years are highlighted with colours. The study year, 2024, is shown with a dashed line. The map is adapted from Ittonen *et al*. (2022), originally published under a CC BY 4.0 (https://creativecommons.org/licenses/by/4.0/) license.

In August 2024, we collected mated wild females in Sweden from four different areas, denominated populations, that were also used in previous studies of the species (Ittonen *et al*., 2022, 2023, 2025). Two populations were sampled at the northern range margin of the species (Rindö, 59.4°N, 18.4°E; and Katrineholm, 59.0°N, 16.3°E), the other two within the species core range in southern Sweden (Vejbystrand, 56.3°N, 12.8°E; and Hässleholm, 56.2°N, 13.7°E; Fig. 1a).

### 2.2. Diapause induction experiment

We reared *L. megera* on potted *Poa annua* (grown from seeds at Stockholm University) inside mesh cages (described in Ittonen *et al*., [2025]), allowing natural photoperiodic and thermal variation. We placed eggs into these field cages on 5 August (early cohort), 12 August (intermediate cohort), and 19 August (late cohort) 2024. On 18 October 2024, we ended the experiment for all cohorts and assessed whether individuals had entered diapause (still being caterpillars) or developed directly (pupae or adults). We then weighed all diapausing caterpillars on 19-21 October to get a proxy for autumn growth rate.

The eggs used in the experiment were F1 offspring of wild females caught earlier in August and brought into the laboratory in Stockholm for egg laying. Using F1 offspring allowed following natural phenology, and significant maternal effects are unlikely to affect our results because the studied populations experience similar summer temperatures (Ittonen *et al*., 2025) and genetic differences in diapause induction reaction norms were demonstrated by Ittonen *et al*. (2022). When possible, we used offspring of the same females for the different cohorts, but, as many females did not live long enough to sufficiently contribute to all cohorts, we also caught later-flying females (Table S1 shows the number of females used for each cohort and population). With the exception of Rindö for the early cohort (80 eggs), we used 100 eggs per population and cohort, which we mixed and then divided into five groups of 20 eggs to be placed in separate cages. In each cage, we placed the eggs within a mesh basket (of approximately 2 × 2 × 2 cm) in the middle of the potted grass, which allowed newly hatched caterpillars to crawl onto the grass. The grass was changed every time it started to wither or if it was consumed by the caterpillars. The cages were placed in an almost unshaded grassland at Tovetorp Research Station (Nyköping, 58.95°, 17.15°E) in a 6 × 13 cage grid (including extra cages for the additional generation) with 1 meter separating cages. Each population–cohort combination’s five cages stood at random locations within this grid. The grass was watered using a solar-powered dripping irrigation system (SOL-C24 and SOL-C24L, Irrigatia), and we also added fertiliser once a week. The cages were checked daily to identify non-diapausing individuals (pupae or adults) and monitor grass quality to minimize variation in host plant quality.

### 2.3. Winter experiment

The winter experiment followed methods described in Ittonen *et al*. (2025). Briefly, we placed all weighed caterpillars into overwintering plastic cups with a tuft of *P. annua* (initially fresh but never replaced) that through a small hole reached water in another cup below. On 22 October 2024, all cups (covered with mesh preventing escape) were placed in the same grassland where the diapause experiment was conducted, with a plywood board covering them from direct precipitation that could drown the larvae. We scored overwinter survival on 26 March 2025. The method used for the early and intermediate cohorts differed from that used for the late cohort. Early- and intermediate-cohort caterpillars overwintered in 1L cups, with all individuals of a cage sharing one cup, but all late-cohort individuals overwintered in individual 0.5L cups. The individual overwintering of the last cohort allowed connecting individual pre-winter weights to winter survival, which was needed for testing the effect of prewinter mass on winter survival. Sample sizes are shown in Table S2.

### 2.4. Additional generation

We used the non-diapausing individuals to test fitness consequences of producing an additional (third) generation at the northern range margin of *L. megera*. Emerged adults were moved from their field cages to a laboratory at Tovetorp Research Station, where they could mate with individuals from the same population of origin. For mating, we used laboratory cages (50 × 50 × 40 cm) with a transparent roof and net in the front. We covered the bottom of the cages with moistened paper towels and provided a solution of sugar and water, dripped onto flowering plants (genus *Kalanchoe*), for feeding.

We divided the eggs laid by the mated females (10 females for Hässleholm and 1 female for Vejbystrand) into groups of 20 eggs to be placed in field cages and reared using the methods described above for the diapause induction experiment. We had five cages for Hässleholm (placed in the field on 28 September 2024) and one cage for Vejbystrand (placed in the field on 11 October 2024). To simulate a warmer autumn, we additionally allowed a set of eggs from the Hässleholm population to hatch in the laboratory before placing them in the field cages as first instar larvae (on 2 October 2024). The two northern populations (Rindö and Katrineholm) did not produce enough non-diapausing adults for successful mating. On 2 December 2024, we assessed survival and instar, and, as done for the previous generation, larvae were placed in individual 0.5-L cups for overwintering. We scored overwinter survival on 26 March 2025 (see Table S3 for sample sizes and winter survival).

### 2.5. Statistical analysis and software

We analysed our data using linear and generalized (binomial) linear mixed models. For the binary diapause response, the explanatory variables were population of origin, cohort (the time when the eggs were placed in the field), and their interaction. For prewinter larval mass, we used a linear mixed model, with population of origin, cohort, and their interaction as fixed effects. Both models included field cage as a random effect. We removed dead individuals from analyses of diapause induction and larval mass (328 individuals out of 1180 died).

For winter survival, we fitted two separate generalized linear mixed models with population of origin and cohort as explanatory variables and overwintering cup as random effect. In the first model we excluded the late cohort because it was handled differently (with caterpillars in individual cups) during winter. In the second model, we instead included only the late cohort, whose single-caterpillar overwintering cups allowed testing the effect of individual weights on winter survival.

We performed all analyses in R 4.3.1 and RStudio 4.3.0. We fitted models with package lme4 (Bates *et al*., 2015). We used package car (Fox & Weisberg, 2019) for Wald tests (of prewinter larval mass), base R for likelihood ratio tests (of diapause induction and winter survival), and emmeans (Lenth, 2025) for Tukey tests (all pairwise comparisons). We made the plots using ggplot2 (Wickham, 2016).

## 3. Results

### 3.1. Diapause induction experiment

Northern individuals were more likely to enter diapause than southern ones (X^2^ = 39, df = 3, *p* <0.001), but this effect was apparent only for the first cohort (Fig. 2) – nearly all intermediate-cohort individuals and all late-cohort ones entered diapause, regardless of population origin. Tukey tests showed significant differences between northern and southern populations as well as between the two southern populations (Hässleholm–Katrineholm *p* <.0001; Hässleholm–Rindö *p* <.0001; Hässleholm–Vejbystrand *p* = 0.016; Katrineholm–Vejbystrand *p* = 0.012; Rindö–Vejbystrand *p* = 0.018). There was no significant interaction between population of origin and cohort (X^2^ = 1.98, df = 6, *p* = 0.92), which reflects a ceiling effect; diapause incidence reached or approached 100% in intermediate and late cohorts across all populations, leaving little variation with which to detect the interaction. Diapause responses did not differ among populations for the other cohorts, even though eight intermediate-cohort individuals of southern origin (4 from Hässleholm, 4 from Vejbystrand) developed directly.

**Figure 2.**
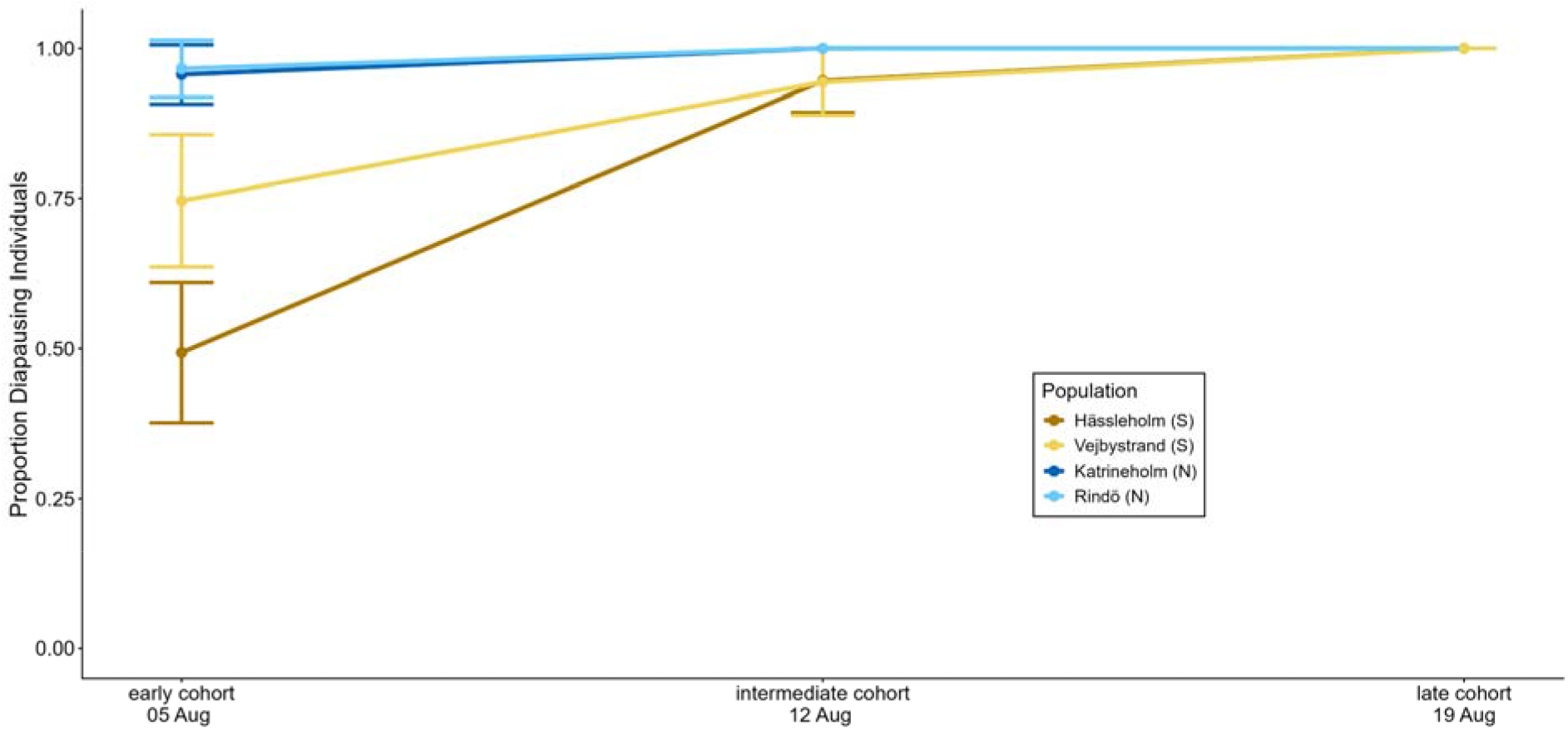
Proportion of diapausing individuals in the three cohorts reared in field cages close to the northern range margin (Tovetorp). See legend for population colours (S = southern population; N = northern population). The bars show 95% confidence intervals (not seen when very close to 1 and thus hidden under the lines).

### 3.2. Larval body mass

By the onset of winter, caterpillars of northern origin had grown larger than those of southern origin, and caterpillars of earlier cohorts were heavier than those of later cohorts (effect of population: X^2^ = 61, df = 3, *p* <0.001; effect of cohort: X^2^ = 91.32, df = 2, *p* <0.001; Fig. 3). All pairwise comparisons, except the one between the two southern populations, showed significant differences (Table S4). There was also a significant interaction between population of origin and cohort (X^2^ = 37, df = 6, *p* = <0.0001). Overall, the north–south differences were more consistent for the two later cohorts, likely reflecting how the longer growing time of the early-cohort allows southern caterpillars to catch up with initially faster-growing northern ones (Ittonen *et al*. 2025).

**Figure 3.**
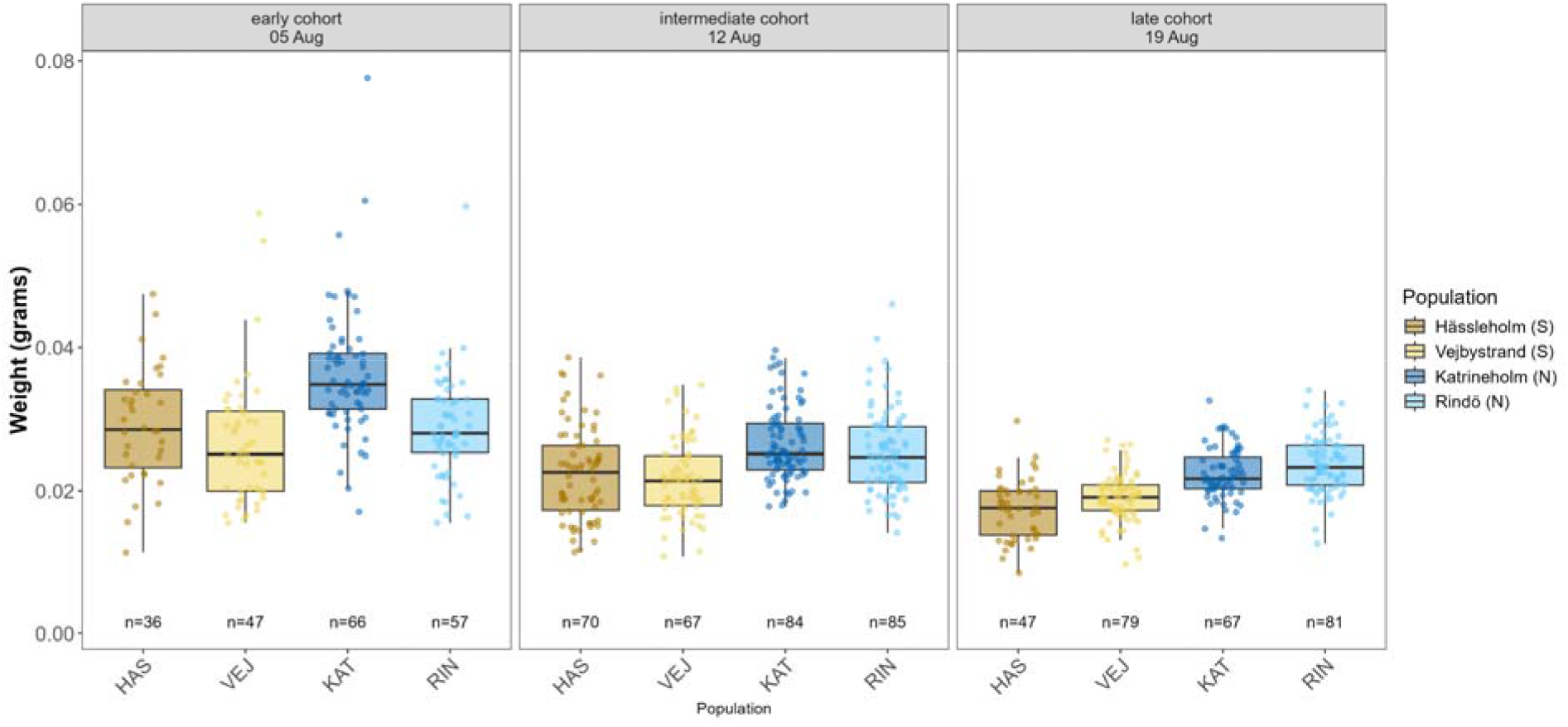
Pre-winter weights of caterpillars per population of origin and cohort. Each panel represents a different cohort, dots show all weighed individuals (sample sizes are shown below the bars), and colours show populations (see legend; S = southern population; N = northern population).

### 3.3. Winter survival

Overwinter survival of early- and intermediate-cohort individuals did not differ among populations (X^2^= 3.70 df = 3, *p* = 0.44) or cohorts (X^2^=0.40, df= 1, *p*= 0.53; Fig. 4a). Winter survival of the late cohort, for which we had individual prewinter mass data, neither differed among populations (X^2^ = 1.72, df=3, *p* = 0.63; Fig. 4a) nor depended on prewinter mass (X^2^ = 1.55, df = 1, *p* = 0.21). Yet, these explanatory variables correlated strongly (Fig. 3), preventing a reliable test of their effects.

**Figure 4.**
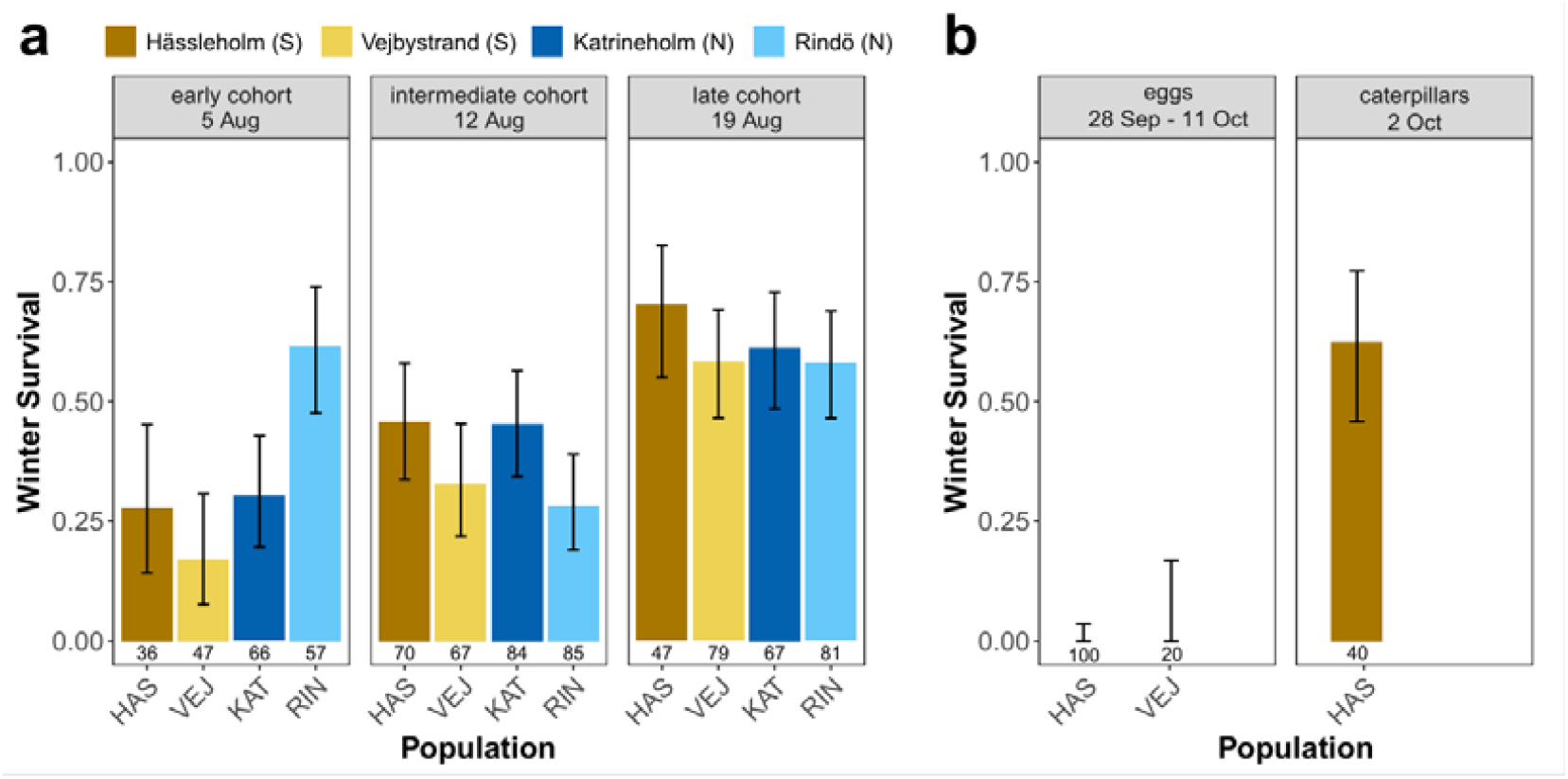
(a) Winter survival per population and cohort of larvae entering diapause in the diapause induction experiment (offspring of second natural summer generation; numbers below the bars=number of diapausing larvae at the start of the winter experiment). Black bars show 95% confidence intervals. Note that late-cohort individuals overwintered individually in small cups, which likely promoted their survival. (b) Winter survival per population and per developmental stage of the offspring of the additional, third, adult generation (non-diapause individuals from the diapause experiment). Black bars show 95% confidence intervals.

Of the additional generation with 100 Hässleholm and 20 Vejbystrand eggs laid by directly developing individuals, none survived winter, although a majority did hatch (Table S3). However, many of the individuals that were first allowed to hatch in the laboratory and then placed in the field as first instar caterpillars entered diapause as second instar caterpillars and survived winter (60%, 24 out of 40 individuals; Fig. 4b).

## 4. Discussion

We reared *Lasiommata megera* from southern core range populations and northern range margin populations in a common garden field experiment close to the species’ northern range margin in Sweden. Individuals of the two northern populations were more likely to enter diapause than southern-population individuals; however, in line with our prediction, this difference was only evident for the earliest of the three weekly cohorts. Moreover, our results clearly demonstrate that direct development into a third adult generation was maladaptive, as directly developing individuals were unable to produce offspring that could enter diapause and survive until spring. Finally, southern and northern populations consistently differed in prewinter body mass, in line with previously demonstrated countergradient evolution (i.e. individuals of northern origin grew faster than individuals of southern origin; Ittonen *et al*., 2025). Still, we found no effect of this size variation on winter survival, and any potential adaptive significance of this pattern is still obscure.

In line with earlier laboratory results showing evolved differences in photoperiodic diapause induction thresholds among the same set of populations (Ittonen *et al*., 2022), caterpillars of northern origin more often entered diapause than individuals of southern origin. Although rapid evolution of photoperiodism for diapause induction has been demonstrated in several other studies (Bradshaw & Holzapfel, 2007; Bean *et al*., 2012; Urbanski *et al*., 2012; Tanaka & Murata, 2016; Batz *et al*., 2020; Reznik *et al*., 2023), the fitness consequences of this genetic variation have rarely been tested in the field (but see Bradshaw *et al*., [2004] for a laboratory demonstration). In our experiment with mostly bivoltine populations, direct development into a third adult generation was clearly a maladaptive choice. Even with the significant benefit of mating and laying eggs in a warm, artificially illuminated laboratory environment, all additional-generation eggs put out into field cages either never hatched or died as first-instar caterpillars (Table S3). This confirms that a southern-type photoperiodic response would, also in this relatively warm year (Swedish Meteorological and Hydrological Institute, 2024), have been maladaptive at the range margin. Nevertheless, when additional-generation eggs were first allowed to hatch in the laboratory and then placed in the field as first-instar caterpillars, a majority of individuals were able to grow and successfully overwintered in the second larval instar. This suggests that a moderately warmer autumn (see Figures S1–2 for logged field temperatures) or laying eggs a week or two earlier could favour the production of a third generation also at the range margin, in particular, if the autumn is warm enough for mating and oviposition. Notably, our first experimental cohort was started on 5 August, while, in the wild, these populations’ earliest second-generation adults are often on the wing at least a week earlier (Fig. 1b).

The phenotypic effect of among-population genetic differences in photoperiodic diapause induction responses depended heavily on when eggs were placed out in the field (Fig. 2). For example, while only half of the early-cohort individuals from the southern population Hässleholm entered diapause, almost all second-cohort individuals – laid only one week later – did enter diapause. This explains the general lack of among-population phenotypic differences in the reciprocal field transplant experiment by Ittonen *et al*., (2025), whose timing conforms with our latest cohort. With that timing, daylength-sensitive larval stages occur in daylengths short enough to induce diapause even in locally maladapted southern individuals (see Table S5 for critical daylengths of each population). Despite this, local adaptation has evolved – most likely because strong selection on just the earliest part of the second annual generation is sufficient – potentially showcasing the trait’s frequently-inferred high evolutionary capacity (Gotthard *et al*., 2026).

Lost generations due to incomplete voltinism shifts have been suggested to play a role in the decline of *L. megera* further south in Europe (Van Dyck *et al*., 2015; Kaiser & Van Dyck, 2025). We confirmed that at least a large part of a third generation would have been lost in the studied year at the range margin. However, this does not necessarily mean that lost generations have large effects on populations. Even in southern Sweden, third generations are always small (Fig. 1b), which limits the potential of lost generations to cause detrimental population-level effects. Furthermore, the survival of individuals put out as first instar caterpillars on 2 October at the northern range margin suggests that southern-Swedish third generations (peaking in mid–late September) could be successful enough to have a neutral or even positive effect on the population. In Belgium, where the lost generation phenomenon has been suggested to harm populations (Van Dyck *et al*. 2015), third generations are large, with potentially large effects on the overwintering population (Kaiser & Van Dyck, 2025), but the growing season is also markedly longer than in Sweden. Therefore, the suggested harmful effects of lost generations in *L. megera* might not necessarily exceed the benefits of reproductive boosts, which appear to counteract decline due to detrimental environmental change in other multivoltine butterflies (Wepprich *et al*. 2025). Successful and lost additional generations are likely to co-occur within and among years, and the net effect of late-season aversion of diapause ultimately depends on reproductive output combined with the offsprings’ winter survival relative to individuals entering diapause in the previous generation. Year-to-year weather variation can shift the balance toward population growth or decline, determining long-term population trends as well as the strength and direction of selection on diapause induction thresholds.

Of the individuals that did enter diapause, caterpillars of northern origin grew faster than caterpillars of southern origin, and earlier-cohort individuals reached bigger sizes than individuals from later cohorts. This pattern of countergradient variation (Conover & Schultz, 1995; Yamahira & Conover, 2002; Carbonell & Stoks, 2020; Kojima *et al*., 2020) closely replicates results from earlier reciprocal field transplant and laboratory common garden studies with the same *L. megera* populations (Ittonen *et al*. 2025). However, neither the earlier study nor ours could establish a clear fitness effect of the resulting variation in prewinter mass. Higher mass is associated with higher winter survival in many other insects (Pullin, 1987; Rozsypal *et al*., 2021; Nielsen *et al*., 2022; Roberts *et al*., 2023), yet we did not find any significant effect of prewinter mass on winter survival. Thus, the significance of faster autumn growth before diapause remains unclear and may depend on the exact ecological circumstances, such as food quality and availability of food during the winter months. Differences in growth rate are seen early on during larval growth (Ittonen *et al*. 2025), so their effects could matter with smaller individuals than those in our experiment, such as those that would hatch even later in August or in September. If so, the voltinism shift occurring further south in Europe could perhaps lead to selection that cancels out the countergradient variation, if the offspring of third generations are pressed to grow fast.

Our common garden field experiment with eggs introduced at three different time points allowed demonstrating the fitness effect of attempting a third generation at the range margin. Moreover, it demonstrated how evolved genetic differences interact with phenology to shape the developmental decisions that caterpillars make. We used well-proven experimental methodology with only small alterations to the natural climate (Ittonen *et al*. 2025) but did provide growing caterpillars with fresh food plants at all times. Still, many of the common and widespread grasses that *L. megera* caterpillars feed on (such as *Poa* sp. *Festuca* sp. and *Dactylis glomerata*) tend to look fresh and healthy in the wild even in late autumn, so food availability would be unlikely to restrict caterpillars in Sweden from developing even in October. Generalizations of our results to other species should nevertheless consider the phenology of host plants and other interacting species. Furthermore, multi-year studies are needed to quantify how interannual climate variability drives the strength of selection on diapause induction and the demographic consequences of partial voltinism shifts. Notably, our results are from a year when a small third generation occurred naturally at the range margin (Fig. 1), probably facilitated by an exceptionally warm early autumn (Swedish Meteorological and Hydrological Institute, 2024; Fig. S1).

We have shown that producing an additional (third) adult generation at the range margin of *L. megera* is a largely maladaptive choice, confirming the adaptive significance of previously demonstrated local evolution (Ittonen *et al*. 2022). The phenotypic effect of genetic differences, however, depends on the exact seasonal timing of the life cycle, and the rapid local adaptation has occurred through natural selection acting only on a limited part of each autumn generation. Our simulation of long-range dispersal from south to north also highlights that local maladaptation to daylength conditions may not severely affect the ability of southern genotypes to colonise northern areas, as a substantial proportion of the new population would survive also in the novel northern conditions. Consequently, this type of dispersal is likely to place a newly established population only somewhat offset from a peak in the adaptive landscape, allowing it to subsequently climb this adaptive peak due to the type of local selection demonstrated here. This may be common since local adaptation of photoperiodism has evolved frequently and rapidly across latitudinal populations of insects (Joschinski & Bonte, 2021; Gotthard *et al*., 2026). Combined, our results suggest that latitudinal differences in daylength are unlikely to restrict climate change–driven range expansions.

## Acknowledgements

We would like to thank the members of the Stockholm University Butterfly Lab for helping to collect the butterflies and for their input on this paper, and Thomas Giegold for his practical assistance at Tovetorp field station. We would also like to thank Katharina Schneider for her help with statistical analyses, figures, and for her support with this paper.

Our work was funded by The Swedish Research Council (to KG; Grant Numbers VR 2017-04159 and VR 2017-04500), Carl Tryggers Stiftelse för Vetenskaplig Forskning (to KG; Grant Number CTS 17:163), and the Bolin Centre for Climate Research (to KG).

