## Supplemental Table S1-S2-S3-S4-S5; Supplemental Figure S1-S2 for "Fitness effects of local seasonal adaptation depend on the timing of reproduction in a range-expanding butterfly"

**Supplementary material**

**Table S1.** Number of families (i.e. wild females) and individuals per family used per population per each cohort in the diapause induction experiment. All population–cohort combinations had five cages with 20 eggs in each, except for Rindö in the early cohort, which had four cages with 20 eggs in each.

| **Cohort** | **Population** | **Number of families** | **Eggs per family** |
| --- | --- | --- | --- |
| Early | Hässleholm | 12 | 8, 8, 8, 9, 8, 8, 8, 8, 8, 8, 8, 8 |
| Early | Vejbystrand | 11 | 10, 9, 9, 9, 9, 9, 9, 9, 9, 9, 9 |
| Early | Rindö | 4 | 22, 34, 21, 3 |
| Early | Katrineholm | 6 | 22, 19, 23, 8, 20, 8 |
| Intermediate | Hässleholm | 6 | 17, 17, 17, 17, 16, 16 |
| Intermediate | Vejbystrand | 6 | 17, 16, 17, 17, 16, 17 |
| Intermediate | Rindö | 7 | 10, 25, 24, 5, 24, 8, 4 |
| Intermediate | Katrineholm | 6 | 20, 19, 5, 22, 19, 15 |
| Late | Hässleholm | 5 | 7, 15, 16, 31, 31 |
| Late | Vejbystrand | 6 | 11, 21, 23, 10, 20, 15 |
| Late | Rindö | 11 | 11, 11, 2, 11, 11, 11, 8, 9, 7, 8, 11 |
| Late | Katrineholm | 6 | 17, 17, 16, 16, 17, 17 |

**Table S2.** Sample sizes for the winter experiment (winter survival) per population and cohort. Early and intermediate cohort individuals overwintered in 1-L cups shared among individuals reared in one cage, while the late-cohort individuals overwintered individually in smaller cups.

| **Cohort** | **Population** | **Sample size** | **Overwintering cup** |
| --- | --- | --- | --- |
| Early | Hässleholm | 36 | 1 L, pooled |
| Early | Vejbystrand | 47 | 1 L, pooled |
| Early | Rindö | 57 | 1 L, pooled |
| Early | Katrineholm | 47 | 1 L, pooled |
| Intermediate | Hässleholm | 70 | 1 L, pooled |
| Intermediate | Vejbystrand | 67 | 1 L, pooled |
| Intermediate | Rindö | 85 | 1 L, pooled |
| Intermediate | Katrineholm | 84 | 1 L, pooled |
| Late | Hässleholm | 47 | 0,5 L, individual |
| Late | Vejbystrand | 79 | 0,5 L, individual |
| Late | Rindö | 81 | 0,5 L, individual |
| Late | Katrineholm | 67 | 0,5 L, individual |

**Table S3.** Survival of the additional generation per population and life stage during autumn and winter. Autumn mortality for the eggs refers to eggs that did not hatch, while winter mortality refers to caterpillars that died during winter. Autumn and winter mortality for individuals put out as caterpillars (life stage) indicate caterpillars that died in autumn and winter, respectively.

| **Life stage** | **Population** | **Starting sample size** | **Autumn mortality** | **Winter mortality** |
| --- | --- | --- | --- | --- |
| Eggs | Hässleholm | 100 | 43 | 57 |
| Caterpillars | Hässleholm | 40 | 7 | 11 |
| Eggs | Vejbystrand | 20 | 10 | 10 |

**Table S4**. Pairwise comparisons of larval prewinter body mass between population and cohort.

| **Cohort** | **Contrast** | **Estimate ± SE** | **df** | **p-value** |
| --- | --- | --- | --- | --- |
| Early | Hässleholm - Katrineholm | -0.24±0.05 | 76.4 | <.0001 |
| Early | Hässleholm - Rindö | 0.002±0.05 | 72.6 | 1.0000 |
| Early | Hässleholm - Vejbystrand | 0.10±0.05 | 100.5 | 0.2710 |
| Early | Katrineholm - Rindö | 0.24±0.04 | 40.5 | <.0001 |
| Early | Katrineholm - Vejbystrand | 0.33±0.05 | 59.2 | <.0001 |
| Early | Rindö - Vejbystrand | 0.10±0.05 | 57.1 | 0.2135 |
| Intermediate | Hässleholm - Katrineholm | -0.20±0.04 | 37.1 | 0.0001 |
| Intermediate | Hässleholm - Rindö | -0.14±0.04 | 36.9 | 0.0048 |
| Intermediate | Hässleholm - Vejbystrand | 0.02±0.04 | 43.9 | 0.9386 |
| Intermediate | Katrineholm - Rindö | 0.05±0.04 | 31.3 | 0.5221 |
| Intermediate | Katrineholm - Vejbystrand | 0.21±0.04 | 38.0 | <.0001 |
| Intermediate | Rindö - Vejbystrand | 0.17±0.04 | 37.7 | 0.0010 |
| Late | Hässleholm - Katrineholm | -0.27±0.05 | 58.5 | <.0001 |
| Late | Hässleholm - Rindö | -0.32±0.04 | 51.9 | <.0001 |
| Late | Hässleholm - Vejbystrand | -0.10±0.04 | 52.8 | 0.1053 |
| Late | Katrineholm - Rindö | -0.06±0.04 | 40.4 | 0.4875 |
| Late | Katrineholm - Vejbystrand | 0.20±0.04 | 41.2 | 0.0012 |
| Late | Rindö - Vejbystrand | 0.22±0.04 | 35.1 | <.0001 |

**Table S5.** The calendar dates at which the actual daylength (including civil twilight) at the experimental site was closest to each population’s critical daylengths (estimated in constant 16 °C conditions by Ittonen *et al*. 2022).

| **Population** | **Critical daylength** | **Date** |
| --- | --- | --- |
| Hässleholm | 16 h 0 min | 26 August |
| Vejbystrand | 15 h 49 min | 28 August |
| Katrineholm | 16 h 28 min | 21 August |
| Rindö | 16 h 35 min | 20 August |

***Temperature logging***

Multiple loggers (HOBO MX Temperature/Light) were used to measure temperatures every 30 minutes, both inside (5 loggers) and outside (5 loggers) the cages where the caterpillars were reared. The loggers were placed inside white plastic tubes (90 mm diameters; 75 mm length), to avoid direct sunlight on the loggers. For the inside loggers the white tubes were attached to the grass’s plastic trays. For the outside loggers the tubes were attached to a wooden stick placed in the ground, to a similar height to the inside logger (about 30 cm from the ground).

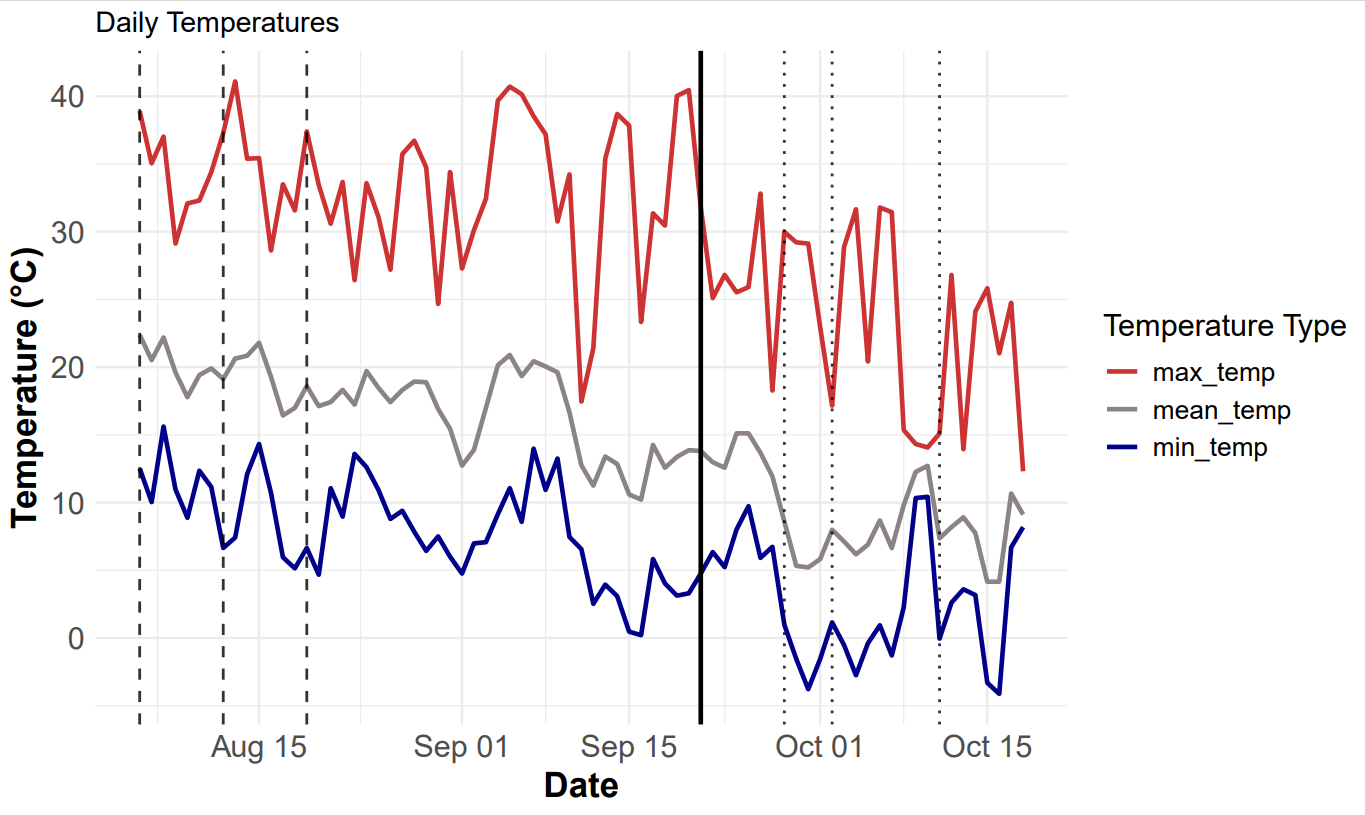

**Figure S1**. Temperature during the diapause induction experiment (5 August 2024 to 18 October 2024), with daily maximum, minimum, and mean temperature shown in red, blue, and grey, respectively. Dashed lines in August show when the eggs were placed in the field for the diapause experiment. The solid line in September shows when the first directly developing adults emerged and mating started in the laboratory, and dotted lines in October show when the resulting offspring (that is, the additional generation) were placed in the field.

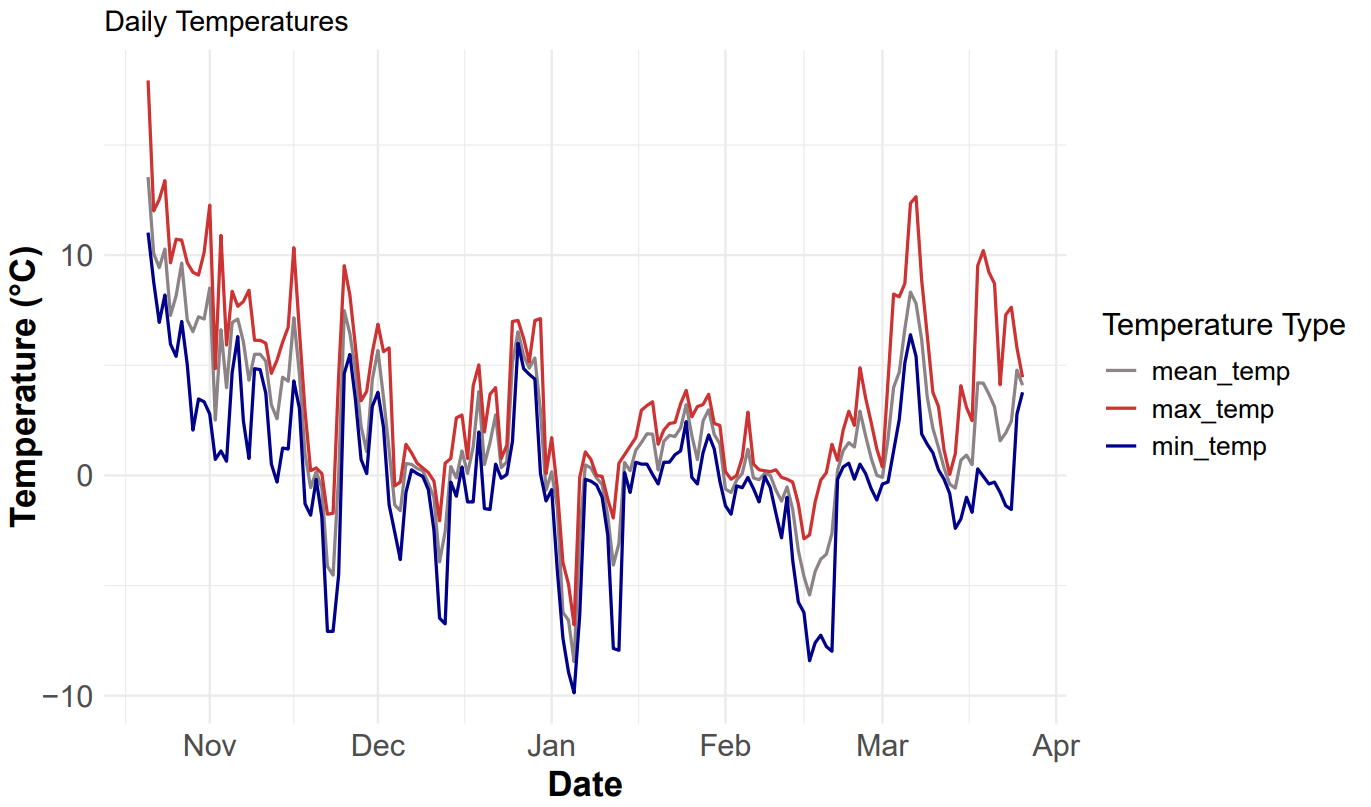

**Figure S2**. Temperature during the winter experiment (22 October 2024 to 26 March 2025), with daily maximum, minimum, and mean temperature shown in red, blue, and grey, respectively.
